# Controllability over stressor decreases responses in key threat-related brain areas

**DOI:** 10.1101/2020.07.11.198762

**Authors:** Chirag Limbachia, Kelly Morrow, Anastasiia Khibovska, Christian Meyer, Srikanth Padmala, Luiz Pessoa

**Affiliations:** Department of Psychology, University of Maryland, College Park, MD, USA; Neuroscience and Cognitive Sciences program, University of Maryland, College Park, MD, USA; Indian Institute of Science, Bangalore, Karnataka, India; Maryland Neuroimaging Center, University of Maryland, College Park, MD, USA; Department of Electrical and Computer Engineering, University of Maryland, College Park, USA

**Author notes:** Corresponding author: Luiz Pessoa, Department of Psychology, 1147 Biology-Psychology Building, University of Maryland, College Park, MD 20742, USA.

## Abstract

Controllability over stressors has major impacts on brain and behavior. In humans, however, the effect of controllability on the responses to stressors themselves is poorly understood. Using functional magnetic resonance imaging, we investigated how controllability altered responses to a shock-plus-sound stressor. Using a between-group yoked design, participants in controlled and uncontrolled groups experienced the same amount of stressor exposure. Employing both Bayesian multilevel analysis targeting regions of interest and standard voxelwise analysis, we found that controllability decreased stressor-related responses across key threat-related regions, notably in the bed nucleus of the stria terminalis (part of the extended amygdala) and the anterior insula. The posterior cingulate cortex, the posterior insula, and possibly the medial frontal gyrus (in exploratory analyses) were uncovered as sites where control over the stressor increased brain responses. Our findings support the idea that the aversiveness of the stressor is reduced when controllable, leading to decreased responses across key regions involved in anxiety-related processing, even at the level of the extended amygdala.

---

Learned helplessness offers a dramatic illustration of the impact of controllability over biologically relevant outcomes (Maier and Seligman 1976). Dogs learned to cross a barrier to avoid a stressor within a few trials. Yet, animals initially given inescapable stressors failed to later learn to escape it. Indeed, research over the past 50 years has revealed that uncontrollable electric shocks in rodents increase defensive reactions to threat including freezing, and subsequently impair instrumental learning in contexts where control is possible (see Maier and Seligman 2016, Moscarello and Hartley 2017). In contrast, controllable stressor exposure is associated with the opposite profile: decreased freezing, increased social exploration, and improved instrumental learning.

At the neural level, the work of Maier and colleagues has uncovered important components of threat controllability, leading to a model in which the ventromedial prefrontal cortex (PFC) plays a central role is blunting the dorsal raphe nucleus’ response to the stressor (Maier 2015). Critically, the serotonergic neurons of the dorsal raphe are only activated if the stressor is uncontrollable. Naturally, these two brain structures do not work in isolation but interface with several others, including the locus coeruleus, the bed nucleus of the stria terminalis (BST), the amygdala, and the periaqueductal gray (PAG). In particular, in the model by Maier and colleagues, the first two areas are conceptualized as inputs and the last two as outputs of the dorsal raphe (Maier 2015).

Controllability research in rodents has inspired a considerable amount of work in humans (Salomons, Johnstone et al. 2004, Salomons, Johnstone et al. 2007, Kerr, McLaren et al. 2012, Collins, Mendelsohn et al. 2014, Hartley, Gorun et al. 2014, Harnett, Wheelock et al. 2015, Montoya, van Honk et al. 2015, Bräscher, Becker et al. 2016, Boeke, Moscarello et al. 2017, Wendt, Löw et al. 2017), some of it focusing on questions related to learning that a stimulus is controllable. For example, participants who learned to actively avoid shocks showed greater responses in the ventromedial PFC (particularly to CS+ or CS-cue offsets) than those who experienced passive omission of shock (Boeke, Moscarello et al. 2017).

However, a key gap in the human literature pertains to the effect of controllability on the responses to the stressors themselves: how do brain regions respond to a stressor event as a function of controllability? In humans, although a few studies speak to this issue (Salomons, Johnstone et al. 2004, Salomons, Johnstone et al. 2007, Harnett, Wheelock et al. 2015, Bräscher, Becker et al. 2016), current knowledge is limited. Critically, how the BST, a key region involved in anxiety-related processing (Davis, Walker et al. 2010) has not been ascertained. Additionally, human studies have treated the amygdala as a unit and not characterized the participation of the functionally distinct basolateral and central amygdala subregions (Amaral, Price et al. 1992, Pessoa 2013). Understanding the roles of the central amygdala and BST is particularly important, as they form part of the so-called “extended amygdala” (Alheid and Heimer 1988, Davis, Walker et al. 2010, Fox, Oler et al. 2015), a functional system that is central to fear- and anxiety-related processing. The PAG’s contribution to pain mechanisms is also central to how the brain processes nociceptive insults (Bandler and Shipley 1994). Are PAG signals modulated by controllability? If so, the finding would provide evidence that brain signals can be affected by controllability even at the level of the midbrain. More generally, extensive debate exists about the extent to which regions such as the amygdala are modulated by high-level factors such as attention (Vuilleumier 2006, Pessoa 2013). Determining how controllability impacts basolateral and central amygdala responses is thus important.

Stressor-related responses are not only confined to subcortical areas. Accordingly, we sought to determine the effect of controllability across cortical regions centrally involved in processing stressors, including the insula and the cingulate cortex. Several theoretical frameworks of anxiety consider the anterior insula to play a central role (Paulus and Stein 2006, Grupe and Nitschke 2013). The anterior/mid-cingulate is important in the appraisal and expression of emotion (Etkin, Egner et al. 2011, Shackman, Salomons et al. 2011). Here, we focused on the anterior mid-cingulate cortex (aMCC), which has been suggested to be an area at the interface between emotion and motor interactions (Vogt and Vogt 2009, Pereira, de Oliveira et al. 2010, Lima Portugal, Alves et al. 2020). We also investigated the ventromedial PFC, which as stated has been implicated in the modulation of responses to stressors.

To examine these questions, we employed the moving-circles paradigm recently developed by our group (Meyer, Padmala et al. 2019), where two circles move around the screen, sometimes moving closer and at times moving away from each other. When the circles touch, participants are delivered a mild electric stressor (Figure 1A; Supplementary Movie 1). Note that circle movement, while smooth, has a high degree of unpredictability. For example, the circles might approach each other such that the stressor is more imminent, then retreat from each other for a period of time. In the present experiment, shock administration was accompanied by a loud buzzer-like sound, increasing the aversiveness of the overall stressor (stressor plus sound). Two separate participant groups were scanned with functional magnetic resonance imaging (fMRI). In the CONTROLLABLE group, when the stressor was administered, its duration could be controlled by pressing a button to turn a virtual wheel (Figure 1B-C). In the UNCONTROLLABLE group, participants pressed a button but it did not have any relationship to stressor termination. In effect, the two groups were yoked such that the aversive stimulation experienced by participants was equated between them.

**Figure 1.**
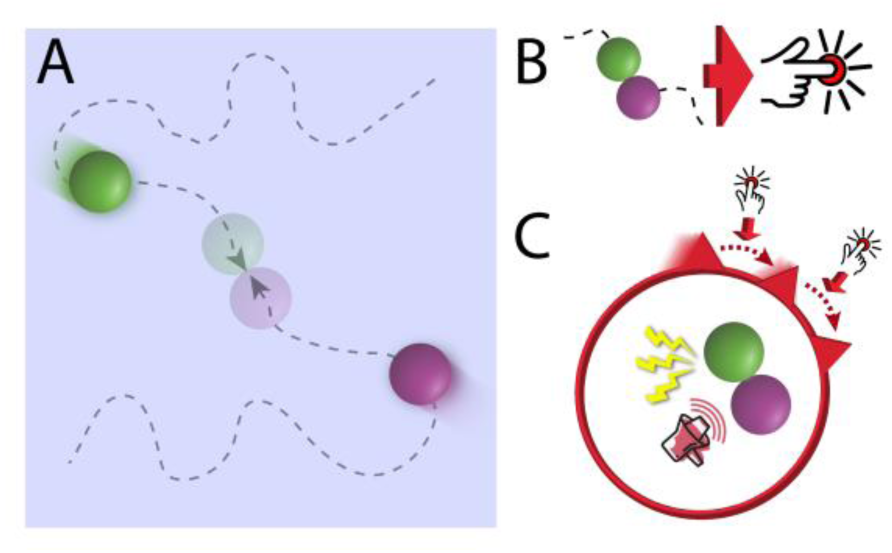
Moving-circles paradigm and controllability. (A) Schematic representation of the display. When the circles collided, a mild shock plus aversive sound was administered. (B-C) For controllable participants, a button press moved the virtual wheel by 1/12 of the circle. The participant had to press it multiple times to terminate the stressor. For uncontrollable participants, the virtual wheel moved the same number of times as that of the yoked participant. Uncontrollable participants were asked to press the button every time the wheel moved; the button press moved the virtual wheel a random amount (not shown). Stressor duration was matched between the two groups.

Whole-brain analysis with fMRI often lacks statistical power to uncover effects at the voxel level, which can lead to poor reproducibility (Cremers, Wager et al. 2017). Therefore, here we sought to focus on a set of regions of interest (ROIs) and leverage the strengths of Bayesian multilevel modeling (BML; Gelman and Hill 2006, McElreath 2020) to estimate the effect of stressor controllability. One of the strengths of BML is that it allows the simultaneous estimation of multiple clustered parameters within a single model (for example, the effects at multiple schools within a district in an educational setting). In the present context, BML allowed the estimation of the effects across multiple ROIs simultaneously (Chen, Xiao et al. 2019). Among the advantages of this approach, information about the effect in one region can be shared across all regions (technically referred to as “partial pooling”). Another important feature is that correction for multiple comparisons is not needed as a single model is estimated.

Given our focus on a targeted set of ROIs and statistical analysis within the context of multilevel modeling, we decided to expand upon the small number of regions mentioned above, while maintaining the set of regions more directly tied to threat-related processing (Figure 2). Accordingly, we considered the anterior hippocampus, which has been discussed in the context of defensive behaviors (Canteras, Resstel et al. 2009, Adhikari, Topiwala et al. 2010, Bach, Guitart-Masip et al. 2014, Kim, Park et al. 2015, Engin, Smith et al. 2016, Qi, Hassabis et al. 2018, Bach, Hoffmann et al. 2019). As stated above, the ventromedial PFC has been implicated in threat processing, in particular controllability. At the same time, posteromedial sites, including the posterior cingulate cortex (PCC) and precuneus have been implicated in certain aspects of threat processing. For example, the PCC was engaged by “slowly attacking” threats (Qi, Hassabis et al. 2018), as well as more distal threat in our moving-circles study (Meyer, Padmala et al. 2019). The PCC/precuneus was more engaged by a distal virtual tarantula (Mobbs, Yu et al. 2010) and distal threat in our moving-circles study (Meyer, Padmala et al. 2019). Accordingly, we added sites along the posteromedial cortex to our set of ROIs so as to investigate the effect of controllability. Finally, we also considered the thalamus, which is increasingly understood to be a key region involved in threat processing (Mobbs, Yu et al. 2010, Salay, Ishiko et al. 2018, Meyer, Padmala et al. 2019).

**Figure 2.**
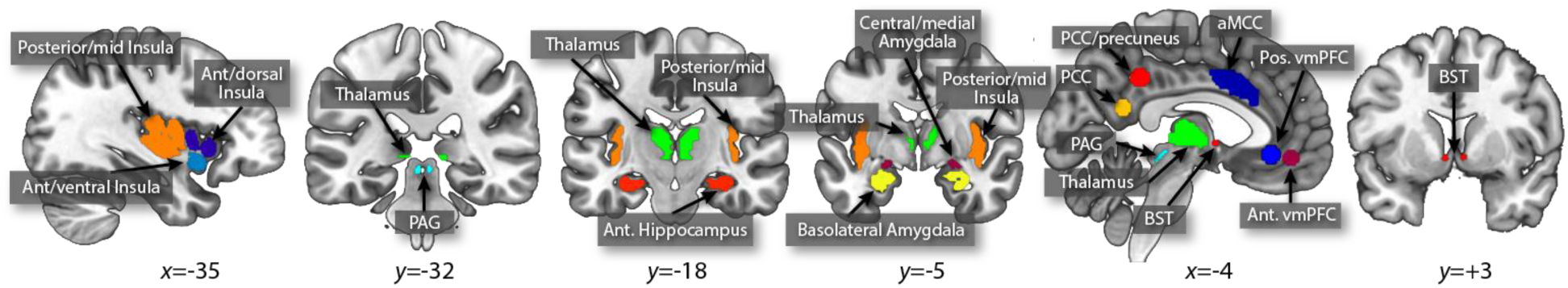
Regions of interest (ROIs). The main analyses targeted a set of brain regions involved in threat processing. ROIs were defined anatomically, with the exception of the thalamus, vmPFC (both sites), PCC, and PCC/precuneus, which were defined based on separate functional datasets. Abbreviations: aMCC, anterior midcingulate cortex; ant., anterior; BST, bed nucleus of the stria terminalis; PAG, periaqueductal gray; vmPFC, ventromedial prefrontal cortex.

## Results

Data were analyzed with Bayesian methods (see **Methods**), including simple tests of behavior, multilevel analysis for a set of 24 target ROIs (one representative time series per ROI obtained by averaging *unsmoothed* functional data to preserve spatial resolution), and multilevel analysis of voxelwise data within the left insula. As Bayesian whole-brain voxelwise analysis was not computationally feasible at present, an additional standard voxelwise analysis was performed. For all analyses, the unit of interest was the yoked participant pair. In all Bayesian analyses, we report evidence for effects in terms of P+, the probability that the effect is greater than zero based on the posterior distribution: values closer to 1 provide evidence that the effect of interest is greater than zero while values closer to zero convey support for a negative effect. We treat Bayesian probability values as providing a continuous amount of support for a given hypothesis; thus not dichotomously as in “significant” vs. “not significant”.

### Behavior

Stressor duration was equated between participant pairs (one participant from each group). Whereas participants from the controlled group executed button presses to terminate the stressor, button presses by uncontrolled-group participants had no bearing on cessation. Controlled participants produced considerably more button presses than uncontrolled participants (173.16 +/- 18.01 vs. 132.27 +/- 55.83 over the experimental session; P+ = 1). Thus, although uncontrolled participants were instructed to press the button with every wheel turn, they did not.

Yoked pairs did not show evidence of difference in state (P+ = 0.49) or trait (P+ = 0.47) anxiety.

### Bayesian multilevel analysis at the level of region of interest

As motivated in the Introduction, to test the effect of controllability, we targeted 24 ROIs (Figure 2). The linear model included covariates for the difference in the number of button presses, and both the average and difference in state/trait anxiety scores. The mean state and trait score for a participant pair tries to capture the fact that some yoked pairs could have high/lower mean scores; the difference in state and trait scores captures the fact that individuals in a yoked pair could be relatively mismatched in terms of their anxiety scores (despite the procedure of matching them during recruitment), but as seen above there was no discernible evidence to support this.

The left and right BST, and the left dorsal anterior insula exhibited very strong evidence for increased responses for uncontrollable participants (P+ of ∼0.99 or higher; Figure 3). A few other ROIs exhibited a moderate amount of support for the same pattern of responses, particularly the left thalamus and the right basolateral amygdala (P+ ∼0.93). Noteworthy evidence for ROIs with controlled greater than uncontrolled responses was not observed (note that in such cases P+ should approach 0).

**Figure 3.**
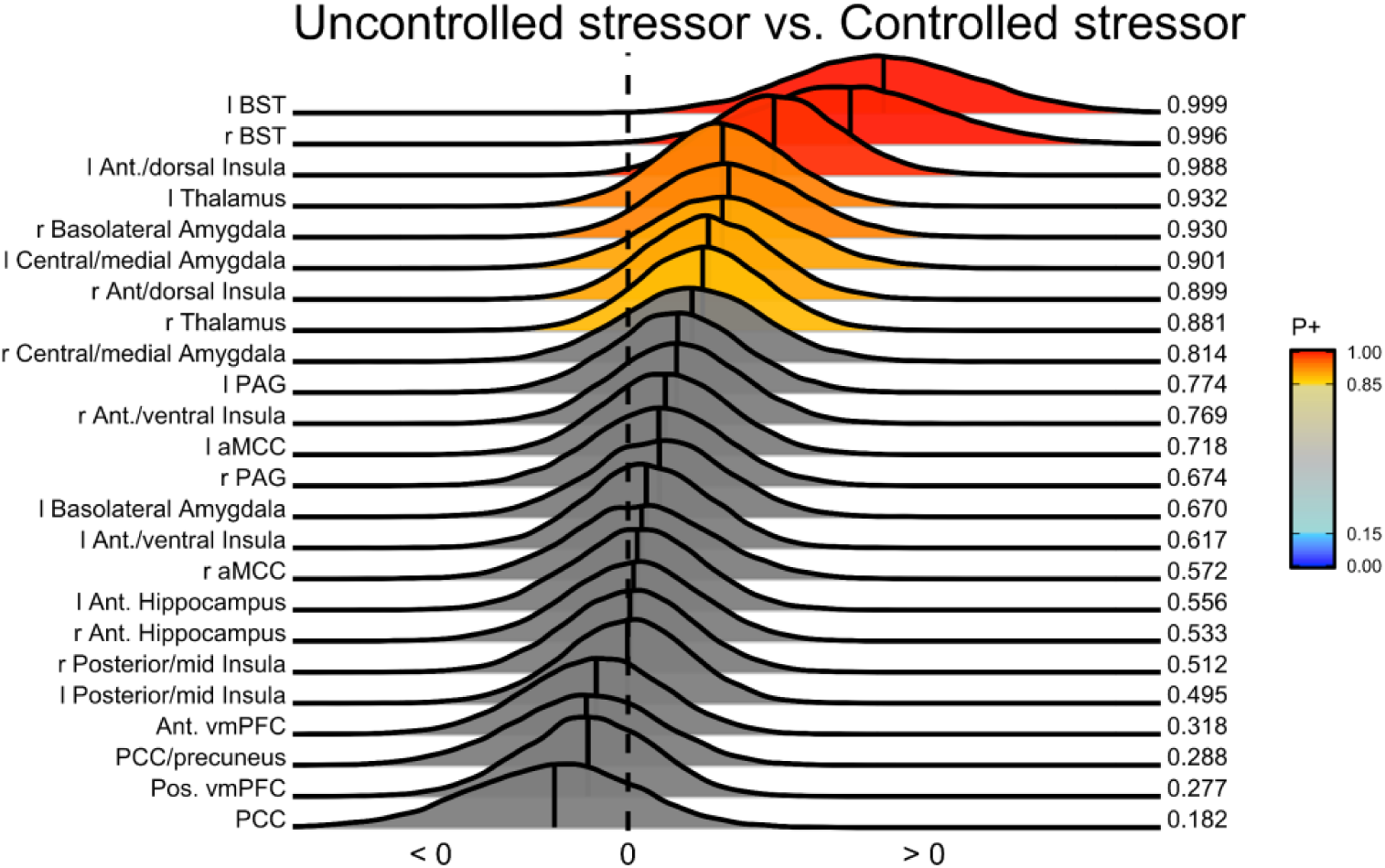
Posterior distributions of the effect of group membership. Three regions of interest exhibited very strong evidence of stronger responses for uncontrolled participants: left/right BST and anterior dorsal insula. P+ values indicate P(effect > 0).

For visualization purposes the responses to the stressor were estimated without assuming a canonical hemodynamic response. Many ROIs generated transient and robust responses to stressor events, such as the BST, PAG, and central/medial amygdala (Figure 4). A relatively weak response was observed in the basolateral amygdala, but the inset shows evidence for a transient response, too. Multiple ROIs along the midline generated negative-going responses, including PCC, precuneus/PCC, ventromedial PFC (two sites), in addition to the hippocampus; the latter exhibited a more bimodal response. These results show that the canonical response provides a good model of responses for most ROIs, with a few exceptions including the PCC, precuneus/PCC, and anterior hippocampus (not because the responses were negative-going but because the shape appears to deviate from the canonical one; for example, for the PCC the negative peak of the response occurred at 2.5 s, which is very early for an event starting at zero).

**Figure 4.**
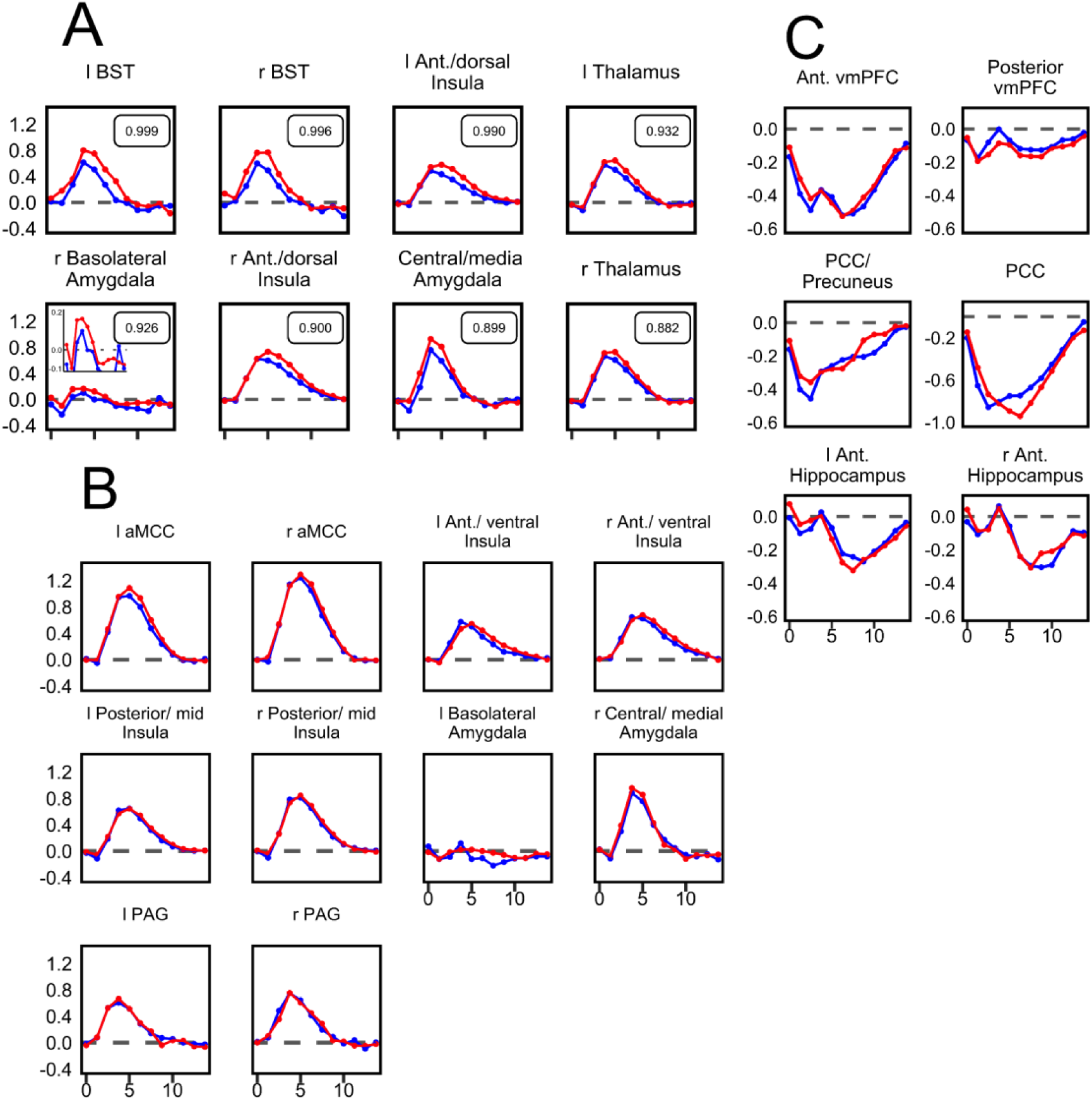
Stressor-related responses (% signal increase), region of interest (ROI) analysis. Unassumed-shape analysis was used to estimate evoked responses to stressors for all ROIs. (A) ROIs with stronger evidence of a controllability effect. P+ values (P(effect > 0)) shown. For the right basolateral amygdala, the inset shows a “zoomed in” version of the response. (B-C) Responses for the remaining ROIs, with the responses to the stressor that were positive-going in (B) and those that were negative-going in (C).

### Individual differences

Evidence for individual differences in state/trait anxiety was limited (Supplementary Figure 1). For example, for the left BST, the contribution of state anxiety difference was positive (P+ = 0.891).

### Brain-skin conductance correlation

We investigated how the trial-by-trial relationship between skin conductance responses (SCRs) and brain responses varied as a function of controllability. We investigated this relationship for the left and right BST, which had the strongest effect of controllability. We initially determined trial-by-trial responses to individual stressors, both in terms of skin conductance and fMRI signals. Trial-based responses were Spearman correlated for each individual, and compared between the two groups (including the covariates). The brain-SCR correlation was higher for uncontrolled compared to controlled participants in the right BST (P+ = 0.963); evidence was only modest for the left BST (P+ = 0.905). We also uncovered potential contributions from state/trait anxiety covariates. For example, in the left BST, the correlation difference (uncontrolled minus controlled) was robustly influenced by the average trait anxiety (P+ = 0.985; Figure 5), such that the higher the average trait anxiety of a participant pair, the higher the correlation difference. Finally, the association was weak for the left dorsal anterior insula.

**Figure 5.**
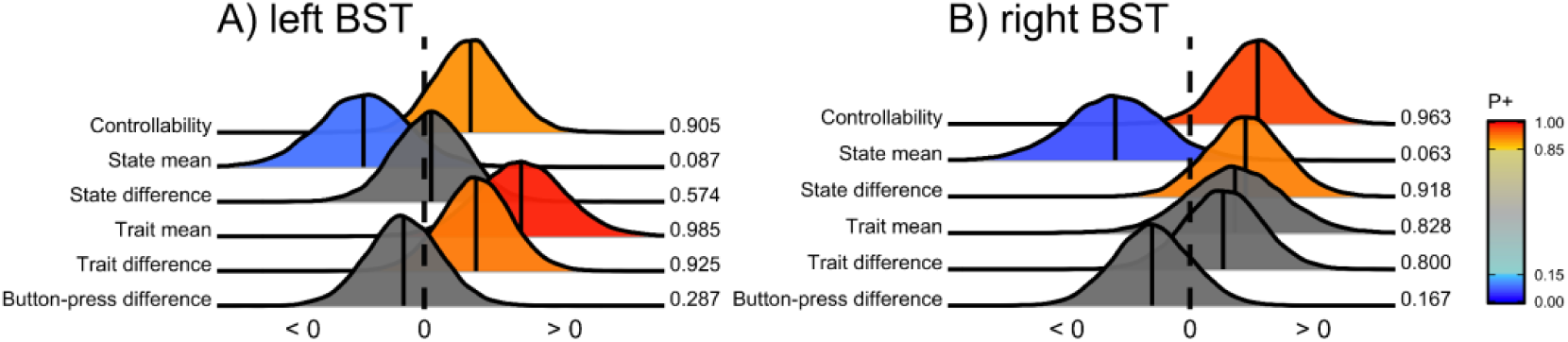
Trial-by-trial brain-skin conductance correlation and controllability. Posterior distributions are shown for the left BST (A), right BST (B). P+ indicates the probability that the effect is > 0.

### Bayesian multilevel analysis of insula voxels

The Bayesian multilevel approach at the ROI level was extended to analyze voxelwise data in the insula (Figure 6). For computational expediency, we analyzed all 941 voxels of the left insula only. Voxels in both the dorsal and ventral anterior parts of the insula exhibited strong evidence for controllable > uncontrollable responses. Notably, we also observed a cluster of voxels in the inferior posterior insula where responses showed a reasonably strong effect in the opposite direction: uncontrollable > controllable (P+ values < 0.05).

**Figure 6.**
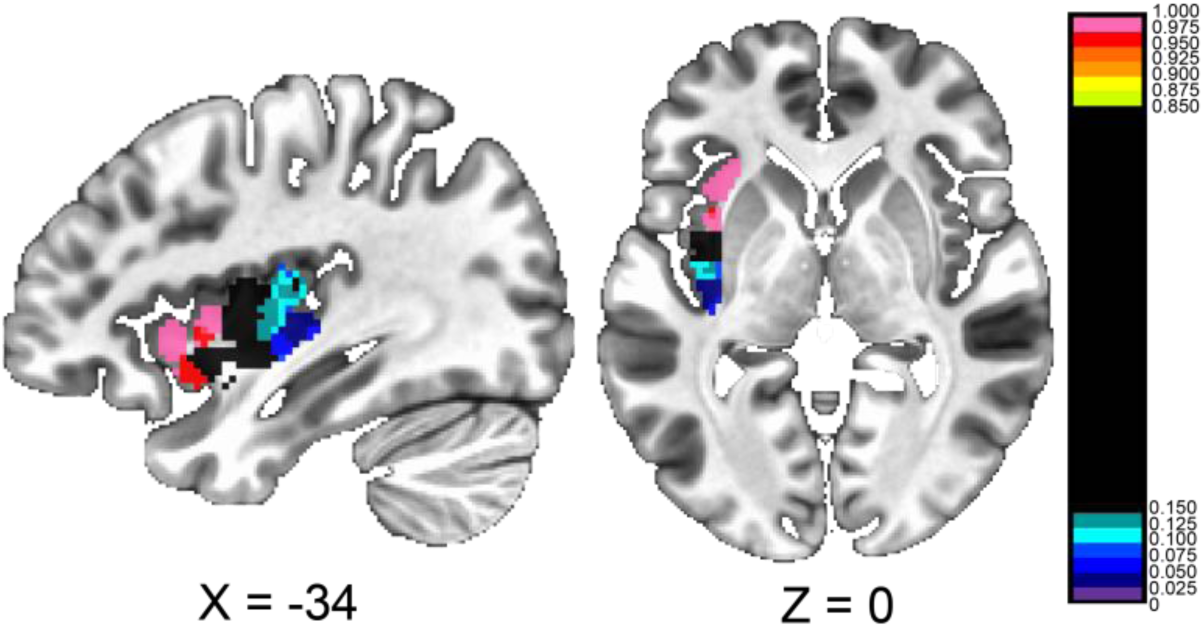
Bayesian multilevel voxelwise analysis of the left insula. A single model analyzed all 941 voxels simultaneously. Thus, no correction for multiple comparisons is needed. P+ indicates the probability that the effect is > 0.

### Whole-brain voxelwise analysis

For completeness, we also ran an additional voxelwise analysis (standard, not Bayesian). Activation clusters were detected in the anterior insula, caudate, and putamen where responses were greater for controlled vs. uncontrolled (Figure 7; Supplementary Table 1; see also Supplementary Figures 2-3). The posterior cingulate cortex was the only cluster that exhibited a response that was greater for controlled participants; at this site, the responses were negative going (Figure 8).

**Figure 7.**
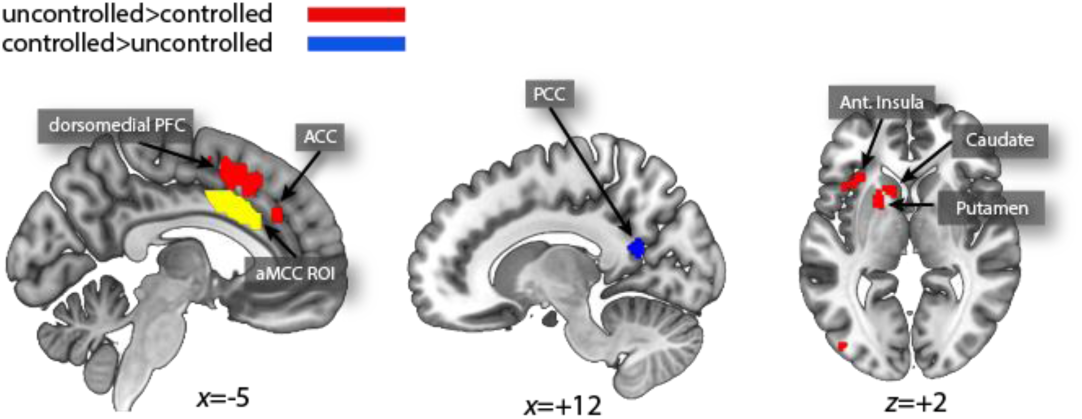
Voxelwise contrast of uncontrolled vs. controlled participants. Maps were thresholded at 0.001 at the voxel level and 0.05 at the cluster level (13 voxels). The yellow region is the anterior midcingulate cortex region of interest shown so that its location can be compared to the voxelwise activation clusters. ACC, anterior cingulate cortex; Ant, anterior; Mid, middle; PCC, posterior cingulate cortex.

**Figure 8.**
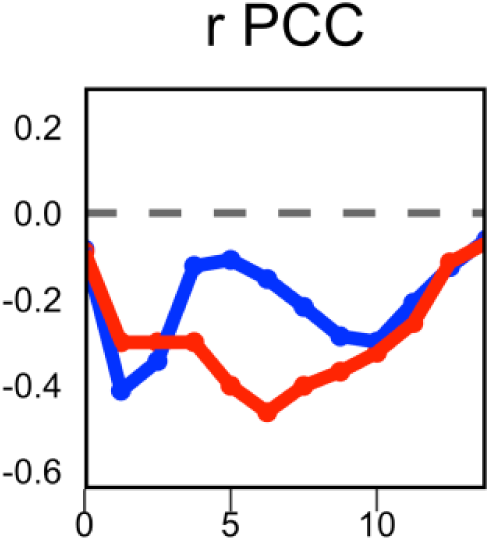
Stressor-related responses (% signal increase), voxelwise analysis. Unassumed-shape analysis was used to estimate evoked responses to stressors for the right posterior cingulate cortex (r PCC); see Figure 7.

As the ventromedial PFC as implicated in controllability in the past ((see Moscarello and Hartley 2017)), in an exploratory fashion, we relaxed our initial thresholding procedure and adopted a voxelwise cluster of 0.005 with a minimum cluster extent of 10 voxels. The contrast revealed additional controlled > uncontrolled clusters (Supplementary Table 2), including the medial frontal gyrus (Figure 9).

**Figure 9.**
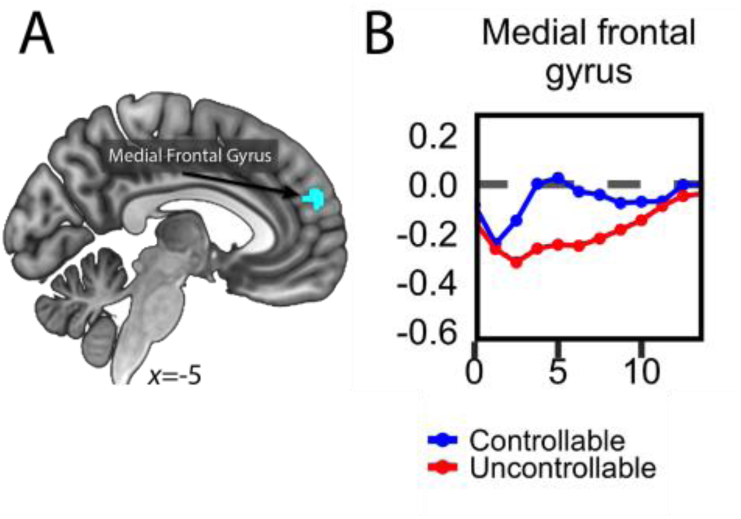
Exploratory voxelwise analysis. (A) Voxelwise threshold at 0.005 and cluster extent of at least 10 voxels: controlled > uncontrolled. (B) Estimated responses based on unassumed shape analysis (% signal increase).

## Discussion

In the present study, we investigated the effect of controllability on evoked responses to shock-plus-sound stressors in humans. Two circles moved around the screen and when they collided the aversive stimulus was administered. Participants in the controlled group could terminate the stressor by actively pressing a button; in contrast, in the uncontrolled group button pressing had no relationship to stressor termination. The experiment was designed by yoking the two groups such that participants experienced the same amount of aversive stimulation. Based on Bayesian multilevel analysis, we investigated responses in a targeted set of ROIs involved in threat processing centered on the BST, basolateral and central amygdala, and PAG, in addition to other brain areas involves in threat processing. The most robust impact of controllability was observed in the BST and the left dorsal anterior insula, where responses were smaller for participants in the controlled group.

Research with both rodents and humans shows that the BST is critically important for the processing of temporally extended and uncertain threat (Davis, Walker et al. 2010, Fox, Oler et al. 2015). In humans, one study showed that responses increased parametrically with threat (a virtual tarantula) proximity. Although data processing employed considerable spatial smoothing, the parametric effect appeared to include the BST. In our previous study using the moving-circles paradigm (where stressor delivery was uncontrollable), we also detected a parametric effect of threat proximity on BST responses; thus, as the circles approached each other activity increased (Meyer, Padmala et al. 2019). This latter study is part of a growing literature in humans that has steadily improved imaging parameters and procedures to image this technically challenging region (e.g., Clauss, Avery et al. 2019, Torrisi, Alvarez et al. 2019, Hur, Smith et al. 2020).

In the present study, the BST exhibited strong transient response to the stressor itself. Critically, the stressor response differed across participant group, being reduced for those having control over the stressor. How can these results be interpreted? The results from the tarantula study (Mobbs, Yu et al. 2010), as well as the threat-proximity effect in the previous moving-circles study (Meyer, Padmala et al. 2019), indicate that more imminent stressors engage the BST more strongly. In addition, in the tarantula study, participants’ “experienced fear” ratings increased substantially when the tarantula was perceived as near. Thus, it is reasonable to assume that stressors perceived to be more powerful are associated with enhanced BST responses. Accordingly, in the current study, the decreased responses in the BST could reflect the diminished aversiveness or perceived impact of the stressor in controllable participants.

The left dorsal anterior insula produced a pattern of responses that paralleled those in the BST. Although the anterior insula is very functionally diverse (Menon and Uddin 2010), it has a prominent role in threat processing, especially when uncertainty is involved (Paulus and Stein 2006, Grupe and Nitschke 2013). The anterior insula is often conceptualized in terms of dorsal and ventral sectors, with the former more closely tied to cognition and the latter with emotion (Deen, Pitskel et al. 2011). Here, only the dorsal sector exhibited a strong effect of controllability. We have argued elsewhere that, although insula sectors exhibit some functional preferences, all parts exhibit very diverse functional profiles (Anderson, Kinnison et al. 2013). We note that our ROI results did not provide support for a controllability effect in the posterior insula. This finding is relevant given the role of this region in interoceptive and pain-related processes (Craig 2002) (but see below).

Our analyses also uncovered a relationship between trial-by-trial brain responses and SCRs. Specifically, the correlation between trial-by-trial responses was higher for uncontrolled participants in the right BST. Although only mild support was observed for the left BST, this ROI showed a robust effect of individual differences. Participant pairs with higher state anxiety exhibited a higher BST-SCR correlation difference (uncontrolled minus controlled). Combined, these results provide support for a close relationship between stressor-related activation in the BST and autonomic responses indexed via skin conductance, as well as the impact of individual differences on this relationship.

We suggest that one of the advantages of the Bayesian analysis performed here is that the results can be interpreted not in terms of a binary decision (significant vs. not significant) but in a graded fashion. Accordingly, the evidence in support of a controllability effect was moderate for the right basolateral amygdala and left thalamus, for example. These regions are, of course, extremely important for threat processing. Furthermore, the present results did not find much supporting evidence for an effect of controllability in the PAG, a key site for threat and pain processing. Finally, our study also adds to the body of data on the “extended amygdala” (BST and central amygdala) in humans (Fox, Oler et al. 2015, Shackman and Fox 2016). Whereas evidence for a controllability effect in the left/right BST was very strong, it was moderate/weak for the left central amygdala and rather weak for the right central amygdala.

As stated in the Introduction, a model of controllability centered on the dorsal raphe has been proposed by Meier and colleagues (Meier, 2015). They propose that the locus coeruleus and the BST provide inputs to the dorsal raphe, while the amygdala and the PAG are its output targets. Functional MRI data do not provide conclusive information regarding directionality of signal flow. Nevertheless, it is noteworthy that the strongest effect of controllability in reducing responses was observed in the BST, with weaker effects in the amygdala and no convincing support in the PAG. Together, these functional results are not easily reconciled with a model with separate inputs and outputs as proposed by Meier and colleagues.

The voxelwise analysis also revealed several noteworthy findings, including group differences in the caudate and putamen. Whereas the striatum has been investigated extensively in the context of appetitive processing, its participation in aversive processing is less understood but has also been documented (Jensen, McIntosh et al. 2003, Delgado, Li et al. 2008, Robinson, Frank et al. 2010). The voxelwise analysis also revealed an effect of controllability in the dorsomedial PFC, a site engaged by negative affect, pain, and cognitive control, with the latter having the most notable contribution (Shackman, Salomons et al. 2011). Another group effect was detected in the ACC at a site that is more strongly engaged by all these three domains (Shackman, Salomons et al. 2011). Of note, the activation cluster did not overlap the anterior midcingulate (aMCC ROI) as defined by Vogt and colleagues (Vogt and Vogt 2009), which is a cortical territory that is centrally engaged in the appraisal and expression of emotion (Etkin, Egner et al. 2011).

The present study uncovered multiple brain regions with negative-going responses to the stressor, including ventromedial PFC ROIs, PCC, and PCC/precuneus. In studies of threat processing, these regions are more strongly engaged during less anxious states (Mobbs, Yu et al. 2010, Somerville, Whalen et al. 2010, Meyer, Padmala et al. 2019). In both the virtual tarantula and the moving-circles paradigm (Mobbs, Yu et al. 2010, Meyer, Padmala et al. 2019), the ventromedial PFC and the PCC/precuneus exhibited parametric threat effects, such that responses increased for *less* proximal threat; conversely, responses decreased for more proximal threat. In the voxelwise analysis, an activation site in the right PCC showed a group effect, where the decrease was less substantial for controllable participants. In fact, for controllable participants, the initial signal decrease reverses and a positive-going response phase is discernable. Unlike the study by Knight and colleagues (2015), we did not observe differences in the ventromedial PFC as a function of controllability. However, previous studies suggest that the ventromedial PFC is particularly important for learning and/or implementing threat prevention strategies (see Moscarello and Hartley 2017). Taken together, in the present study the PCC was a key site expressing a controllability effect such that control over the stressor increased brain responses.

Further evaluation of voxelwise activation maps in an exploratory fashion uncovered additional clusters with controlled > uncontrolled responses. The medial frontal gyrus cluster is particularly relevant given previous human studies that have uncovered the anterior medial PFC sites that play roles in adaptive coping, for example (Collins, Mendelsohn et al. 2014, Bräscher, Becker et al. 2016, Goodman, Harnett et al. 2018). As these activations were observed during follow-up exploratory analyses, we consider them more tentatively. Nevertheless, it is noteworthy that the medial frontal cluster included 30 voxels, and at a 0.005 voxelwise threshold clusters of 27 or more voxels are significant at the cluster level of 0.05. The challenges of corrections for multiple comparisons are considerable in neuroimaging, and intense debate and method development has occurred in the past few years (Eklund, Nichols et al. 2016, Li, Nickerson et al. 2017, Cox 2019). In this regard, we discuss the motivation behind Bayesian multilevel modeling in (Chen, Taylor et al. 2020).

The Bayesian voxelwise analysis of the left insula demonstrated how our modeling approach can be effectively extended to voxel-level data. We found a cluster of voxels in the posterior insula where controlled responses were greater than uncontrolled. These results are noteworthy given the finding that Christianson and colleagues (Christianson, Jennings et al. 2011) found evidence for controllability-related “safety signals” in this area’s potential role in dampening BST responses during controllable conditions.

Is it possible that some of our results were driven by differences in button presses? The data were analyzed with a model that included a parametric regressor for the number of button presses at the participant level, and a covariate for the difference of button presses (between the yoked participants) at the group level. In addition, all effects of controllability were such that responses were greater for the uncontrollable group, which was the one that pressed the button *less* often. Three notable sites exhibited higher responses for controllable vs. uncontrollable participants: right PCC (standard voxelwise analysis), left posterior insula (Bayesian voxelwise analysis of the insula), and medial frontal gyrus (exploratory standard voxelwise analysis). At the voxelwise level, we did not detect any activation that was sensitive to the number of button presses at the corrected statistical threshold. Nevertheless, we inspected the results at minimal threshold levels. At an uncorrected level of 0.01 at the voxel level with five contiguous voxels, we observed only one region that overlapped the activations of Table 1, namely the left anterior intraparietal sulcus (Supplementary Figure 3). Therefore, it is unlikely that the responses in the right PCC, left posterior insula, or medial frontal gyrus were unduly influenced by button presses.

In summary, in the present paper we investigated how controllability alters the processing of a shock-plus-sound stressor in the human brain. Using a between-group yoked design, participants in the controlled condition were able to press a button to turn a virtual wheel to stop the stressor. Although stressor delivery was identical for the two groups, most notably controllability had a robust impact on BST and anterior insula responses, such that they decreased across an extended set of brain sites. Our Bayesian multilevel modeling approach also revealed moderate evidence for decreases in the central amygdala and thalamus. When combined with other studies of threat processing, our findings support the idea that aversiveness of the stressor is reduced when it is controllable leading to decreased responses in key anxiety-related brain regions. Finally, we found three regions that exhibited greater responses in controllable participants: the posterior cingulate cortex, the posterior insula, and the medial frontal gyrus. Thus, in addition to the ventromedial PFC, which has been identified as key region engaged when conditions are controllable, the sites observed here deserve further studies in humans.

## Methods

### Procedure and Stimuli

Two circles moved around on the screen in a smooth fashion. When they collided with each other, an unpleasant mild electric stressor was delivered together with an aversive sound. The algorithm to generate the motion of the circles generated approach and retreat periods of varying durations (2-9 seconds), but with most of them lasting more than 6 seconds of approach or retreat. The total amount of time during which the circles were approaching or retreating was equated. Overall, circle motion was designed to be smooth but still relatively unpredictable in terms of the pattern of approach/retreat. In particular, multiple instances of “near misses” were incorporated such that the circles retreated once they were close to each other.

Visual stimuli were presented using PsychoPy (http://www.psychopy.org/) and viewed on a projection screen via a mirror mounted to the scanner’s head coil. Each participant viewed the same sequence of circle movements. The experiment included 6 runs (8 minutes each), each of which had two 3-minute blocks (2 participants had only 5 runs; 3 participants had only 4 runs; the matching runs were eliminated from the yoked participant’s data). During each block, the circles appeared on the screen and moved around for 180 seconds; blocks were separated by a 30-second off period during which the screen remained blank. A 15-second blank screen was presented at the beginning of each run; a blank screen was presented at the end of the run.

In each block circles collided 0-3 times. A total number of 25 collisions occurred during the course of the experiment (for participants with only 4 and 5 runs, 17 and 20 collisions, respectively). Each collision resulted in the delivery of an electric stressor and of an aversive sound stimulus. The electric stressor (comprised of a series of current pulses at 50 Hz) was delivered by an electric stimulator (Model number E13-22 from Coulbourn Instruments, PA, USA) to the fourth and fifth fingers of the non-dominant left hand via MRI-compatible electrodes. To calibrate the intensity of the stressor, each participant was asked to choose his/her own stimulation level immediately prior to functional imaging, such that the stimulus would be “highly unpleasant but not painful.” After each run, participants were asked about the unpleasantness of the stimulus in order to re-calibrate stressor strength, if needed. Each collision also resulted in the playing of a buzzer-like aversive sound that was on for the duration of the stressor stimulus. The loudness of the sound was set to be “loud but not uncomfortable” by each participant (but never exceeded 85 dB).

Skin conductance response (SCR) data were collected using the MP-150 system (BIOPAC Systems, Inc., CA, USA) at a sampling rate of 250 Hz by using MRI compatible electrodes attached to the index and middle fingers of the non-dominant left hand. Due to technical problems SCR data were not available for 4 participants (the corresponding data for the yoked participants were also not utilized).

### Virtual wheel procedure and participant yoking

When a stressor was delivered, the screen turned white and a red dial appeared around the circles (Figure 1C). Subjects in the controllable group were told that moving the wheel allowed them to terminate the stressor. The wheel turned clockwise by 30 degrees (1/12 of the circle) with every button press. The number of rotations required to terminate the stressor started at one and increased after each successful stressor termination up to 12 total button presses. This procedure was implemented so as to parallel the rodent procedure adopted by Maier and colleagues (for example, Amat, Paul et al. 2006). The procedure had the effect of increasing the stressor duration as the experiment progressed. If a subject failed to escape a stressor in a run, data from that run was not utilized (one participant had 1 run discarded).

Subjects in the uncontrollable group experienced the exact same duration of stressor delivery as yoked participants in the controllable group. Subjects in the uncontrollable group saw the dial rotate on its own during stressor delivery. The timing of the rotation of the dial was matched to that of the yoked controlled participant, but the amount of rotation was random to avoid any sense of progression. Participants were told to press the button whenever they saw the dial rotate, so as to maintain the number of button presses closely matched to that of the yoked controllable participants.

### Regions of interest

We focused on the following ROIs, which are shown in Figure 2. Subcortically, we targeted the following regions: Amygdala (2 sectors), BST, PAG, and thalamus. Cortically, we targeted the following regions: Insula (3 sectors), anterior midcingulate cortex, PCC, PCC/precuneus, and ventromedial PFC (2 sites).

The amygdala ROIs were defined based on the anatomical masks by Nacewicz (2014), which we used in an earlier version of the moving-circles paradigm (Najafi, Kinnison et al. 2017). We created amygdala ROIs for the central/medial amygdala (the central amygdala is too small to be separated from the medial amygdala with the present resolution) and for the basolateral amygdala (union of BLBM and La masks).

The PAG ROI was defined anatomically based on the definition by Erza et al. (2015). However, as multiple voxels from the original mask are adjacent to the cerebrospinal fluid aqueduct, we eroded the mask to exclude these voxels, as follows. Using a separate fMRI study (Meyer, Padmala et al. 2019), we evaluated the signal quality of aqueduct and non-aqueduct voxels. The aqueduct contained no voxels with a signal-to-noise ratio (SNR) greater than 20, whereas SNR within the original Ezra mask spanned from 16 to ∼40. To exclude aqueduct-containing voxels, we eliminated those with SNR less than 25, effectively staying clear of the aqueduct.

The BST ROI was defined anatomically according to the probabilistic mask of the BST (at 25% threshold) reported by Blackford and colleagues (Theiss, Ridgewell et al. 2017), as used in our previous study (Meyer, Padmala et al. 2019).

The anterior hippocampus was defined based on the Freesurfer (https://surfer.nmr.mgh.harvard.edu/) parcellation of the hippocampus. Only the component anterior to a y of −20 (MNI coordinates) was retained.

The insula ROIs were defined based on the masks by Faillenot et al. (2017): dorsal anterior insula (union of the anterior short gyrus and the middle short gyrus), ventral anterior insula (anterior inferior cortex), and middle plus posterior insula (union of the posterior long gyrus, anterior long gyrus, and posterior short gyrus).

The anterior midcingulate cortex was defined based on the mask from Destrieux et al. (2010). Additional cingulate/midline ROIs and medial PFC ROIs were defined based on functional data from separate studies. In all cases they were spherical ROIs with 5 mm radius. A ventromedial PFC ROI (called posterior ventromedial PFC) was defined based on the coordinates provided by Boeke et al. (2017). A second ventromedial PFC ROI (called anterior ventromedial PFC), as well as PCC and PCC/precuneus ROIs, were based on functional data from our previous moving-circles study (Meyer, Padmala et al. 2019), specifically, by evaluating the effect of threat proximity (contrast of farther vs. closer circles); in these three cases, activation was stronger when the circles were farther (the opposite of a region such as the anterior insula).

## Statistical analysis

Statistical analysis was performed with a two-level procedure: estimation of regression coefficients at the individual level, followed by group analysis. Participant-level estimation adopted a standard multiple regression approach. The main group-level analysis followed a Bayesian multilevel modeling framework (Gelman and Hill 2006, McElreath 2020), which has been developed recently for fMRI data (Chen, Xiao et al. 2019). This analysis was performed at the level of ROIs. For completeness, we also performed a parallel standard voxelwise group-level analysis. As at the moment a voxelwise Bayesian multilevel analyses is not computationally feasible, the group-level analysis compared uncontrolled and controlled groups via a paired t test (given participant yoking).

### Participant level

Preprocessed fMRI data of each participant was analyzed using multiple linear regression implemented in AFNI’s 3dDeconvolve program. The analysis was restricted to gray-matter voxels. In our previous studies using similar versions of the moving-circles paradigm, we observed effects of direction (approach vs. retreat) that varied as a function of circle proximity (defined based on the Euclidean distance between the circles); in other words, an interaction. Accordingly, in the present analysis, we subdivided the entire range of distances between circles into proximity segments: “near” captured periods during which the circles were in relatively close range (>33% of the maximum possible distance); “far” captured the remaining time. Note that, by definition, a stressor, which was the event of interest in our analyses, only occurred when proximity was “near”. Every stressor event varied in duration to some extent (range: 160 to 4300 ms). Accordingly, a regressor was defined based on stressor onset times and their durations.

To account for other contributions to fMRI signals, three additional regressors were considered: direction (+1 during approach, −1 during retreat), speed (temporal difference of proximity values), and the interaction between direction and speed (mean centered). Direction and speed regressors were set to zero during stressor administration. These three regressors were defined for the “near” period and, separately, for the “far” period. Given our interest in characterizing stressor-related responses, here we were only interested in the “near” period. Note that a stressor event always followed an approach period, but the relationship between approach segments and the stressor was variable (see also collinearity below). In particular, when the circles approached each other, even in the near space, they frequently altered course and retreated from each other. Of these approach-to-retreat changes, multiple of them were designed to be “near misses” so that the circles came very close together before retreating.

When the circles touched, the stressor was administered, and participants in each group pressed the button (typically multiple times). Although participants were yoked to match stressor duration, the number of button presses could potentially vary between yoked pairs. Accordingly, in addition to the stressor regressor described above, a second regressor was employed that modeled a participant’s number of button presses. This parametric regressor had the same onset and duration as the basic stressor regressor, but was mean centered around the mean number of button-presses to reduce collinearity (mean centering was done on a run-by-run basis because the required number of button-presses increased along the session). Overall, the parametric regressor captured signal variance that was linked to the number of button presses, thereby minimizing potential differential contributions of the participant’s overt behavior.

All the above regressors were convolved with a standard hemodynamic response as defined by the gamma variate model (Cohen 1997). In addition, six motion parameters (three translational and three rotational) and their discrete temporal derivatives were included in the model to account for head motion. To account for baseline and drift in the MR signal, linear and non-linear polynomial terms (up to forth order) were also included in the model. We evaluated collinearity and computed the variance inflation factor associated with all variables (James, Witten et al. 2013). The maximum value was 1.9, revealing that regressor correlation did not unduly compromise parameter estimation. In particular, the parametric variable based on the number of button presses did not lead to high regressor correlation given that it was mean centered (the correlation of this regressor with others was < 0.5). Note that although suggested variance inflation factor cutoffs are necessarily crude, a value of 10 is often described as reason for concern; more conservative suggestions consider a value of 2.5 or higher of possible concern (O’Brien 2007). However, as discussed Mumford et al. (2015), collinearity is not a major concern in two-level statistical analyses such as with fMRI data.

### Bayesian multilevel statistical analysis: Region of Interest

The null hypothesis significance testing (NHST) framework has come under increased scrutiny in recent years. In particular, the hard threshold of 0.05 has come under attack, with reasonable researchers calling for stricter thresholds (Benjamin, Berger et al. 2018) or, conversely, for the dichotomous use of p-values to be abandoned (McShane, Gal et al. 2019). Our own approach, which is described in more detail elsewhere (Chen, Taylor et al. 2020), does not consider a binary threshold (“significant” vs. “not significant”) to be a productive way to evaluate statistical evidence. Accordingly, in the present paper, wherever possible, we employed Bayesian statistical analysis (all analyses except the voxelwise case).

The Bayesian framework attempts to estimate the probability of a research hypothesis H given the data, P(H | data). The framework is not typically formulated to generate a binary decision (e.g., “real effect” vs. “noise”) but instead to obtain the entire probability density distribution associated with P(θ | data), where θ is the parameter being estimated. Such posterior distribution allows the quantification of, for example, P(θ > 0 | data), the area under the curve above 0 which we call P+. Values closer to 1 provide evidence that the effect of interest (e.g., mean, difference of means, etc.) is greater than zero (conditional on the data, the prior distribution, and the model); values of P+ closer to zero convey support for a negative effect (for example, P+ = 0.01 indicates that the probability of the effect being positive is only 0.01, which implies that the probability of it being negative is 0.99).

The posterior distribution provides a summary description of the likelihood of observing parameter values given the data, so it naturally conveys variability. Some authors use cut-off points to summarize “strong”, “moderate”, or “weak” evidence, but we encourage an approach that both quantifies and qualifies the evidence, without making decisions in terms of “passes threshold” versus “fails to pass threshold”. Note that we do not employ Bayes factors, which some have advocated as a potential feature of Bayesian modeling. Because Bayes factors consider the probability of “null” effects (e.g., a mean of zero) versus an “alternative” effect (e.g., a mean different from zero), we believe its use is often problematic, because formulating the problem in terms of “null” effects of exactly zero is often unrealistic (because effects of experimental manipulations are seldom zero), thus largely inflating the evidence for the alternative hypothesis (creating “large” Bayes factors). See Chen et al. (2020) for further discussion. Finally, given that we do not view thresholding as adequate, the P+ probability values that we provide are (by definition) “one-sided”. For readers who absolutely insist on comparing P+ values with standard cut-offs that are “two-sided”, they should bear in mind our definition.

We employed a Bayesian multilevel analysis at the level of ROIs (Chen, Xiao et al. 2019). ROI-level data employed the average-across-voxels time series for each ROI; thus, 24 representative time series were employed. Averaging was performed on *non*-spatially smoothed data to avoid degradation of spatial resolution. In the approach adopted, the data from all ROIs are included in a single multilevel model that evaluates the effects of interest. By doing so, the contributions to fMRI signals of subject-level effects (i.e., subject effect across conditions), and ROI-level effects (i.e., ROI effect across subjects), can be accounted for in a model that simultaneously ascertains the effect of controllability. The “output” of the Bayesian multilevel model comprises only one overall posterior that is a joint distribution in a high-dimensional parameter space; thus, no correction for multiple comparisons is needed (Gelman, Hill et al. 2012). For summary purposes, posteriors of the effects for every ROI can be plotted separately; but they are not independent and technically are simply marginal distributions (that is, projections along particular variables). For formal details of the approach adopted here, please refer to Chen et al. (2019); for a less technical exposition, see Chen et al. (2020). In contrast, based on the standard testing approach, the effect at each region is estimated independently from that at other regions, which calls for a correction for multiple comparisons.

The central question of interest was to compare the difference in stressor responses between the two groups:

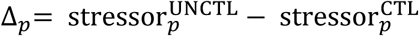

where the superscript UNCTL and CTL refer to the two groups, *p* indexes paired/yoked participants, and `stressor` refers to the parameter estimates from multiple regression from the first-level analysis. However, the difference between the two groups could be itself influenced by factors that differentially affected them, including differences in state and trait anxiety between yoked participants. In addition, differences in button-presses could also influence group differences. Thus, we considered these three types of difference as covariates to be included in the model. Note that the button-press contribution was probably negligible because the estimation of participant-level responses included a parametric regressor for button-presses. In addition to differences in state/trait scores, their sum (or average) could also be a factor. To see this, consider that a yoked pair with higher average state anxiety could potentially produce larger differential stressor-related responses than a pair with lower average state anxiety. Accordingly, we included average state and trait anxiety scores as covariates in the model to evaluate how the effect of controllability was affected by them. Thus far, the model can be summarized as follows:

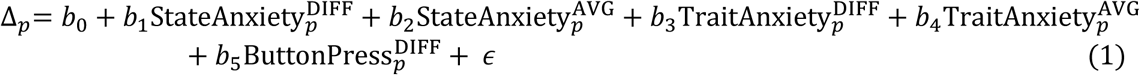

where p indicates a pair of yoked participants, `diff` indicates the difference score, and `avg` indicates the mean score. In our sample, the correlation between state and trait anxiety was 0.5; between state average and state difference was 0.05; and between trait average and trait difference was −0.22.

Within a multilevel framework, we extended the model above to simultaneously consider the contributions of both participant and ROI effects. This allowed us to model a term (intercept) per participant pair (capturing the yoked-pair’s specific contribution relative to the overall intercept), and a term (intercept) per ROI (capturing the ROI-specific contribution relative to the overall intercept), in addition to an overall intercept term. Likewise, each covariate could be modelled via an overall slope plus a slope that was ROI specific. In the terminology of linear mixed effects models, such “varying effects” models, include both “varying intercepts” and “varying slopes” (Gelman and Hill 2006, McElreath 2020). Although linear mixed effects models can be very powerful, parameter estimation can be problematic and/or not possible. However, a Bayesian formulation seamlessly allows the efficient estimation of parameters, while providing interpretability in terms of evidence. The appendix describes the model formally for completeness, including the weakly-informative priors employed, which had negligible impact on the posterior distributions.

All BML models were implemented using `rstan` (https://mc-stan.org/users/interfaces/rstan) which is the `R` interface to the probabilistic `Stan` language (https://mc-stan.org/). `Stan` estimates posterior distributions with state-of-the-art Monte Carlo Markov Chain methods. Estimation convergence was evaluated with Rhat < 1.1 (all values near 1.0)

### Bayesian multilevel statistical analysis: voxelwise within the insula

Whole-brain voxelwise analysis via BML modeling was computationally prohibitive with current resources. Here, we analyzed 941 voxels of the left insula according to the same multilevel strategy outlined above. As the Bayesian model estimated a single multi-dimensional posterior distribution, no correction for multiple comparisons is needed was needed (Gelman, Hill et al. 2012).

Because the insula was anticipated to be functionally heterogenous, we first subdivided the insula into anatomically defined 3 ROIs: dorsal anterior, ventral anterior, and mid-plus-posterior insula (see supplementary methods). Next, each ROI was subdivided into sub-ROIs containing comparable number of voxels. To do so, we simply clustered voxels based on the *xyz* MNI coordinates (via k-means clustering). In this manner, the insula was subdivided into 11 small sub-ROIs (each containing 75-99 voxels). The multilevel model simultaneously estimated the contributions of the subject, sub-ROI, and voxel.

### Additional Bayesian tests

To evaluate group differences of button presses, we employed the R brms package (Bürkner 2017) in which builds upon Stan (https://mc-stan.org/). For the button-press analysis, a simple intercept model was evaluated:

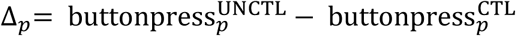

where `buttonpress` refers to the number of button presses (other notation as in equation 1). We assumed a prior distributed as a t distribution which accommodates data skew and outliers well (Kruschke 2014). Analogous models were used to test for differences in state and trait anxiety.

We also evaluated the relationship between trial-by-trial responses in select ROIs (left and right BST, and left dorsal anterior insula) and trial-by-trial responses in skin conductance. Brain-SCR Spearmann correlations were initially computed and Fisher-z transformed. Subsequently, they were tested via a linear model that included state and trait anxiety scores, as well as differences in button presses. In the estimation of the posterior distributions, the effect of controllability was captured by the model’s intercept. Additional slope parameters evaluate the contributions of the covariates. Results were summarized in terms of P+ values, as defined previously.

## Acknowledgments

Research supported by the National Institute of Mental Health (R01 MH071589 and R01 MH112517).

## Supplementary Methods

### Participants

One hundred and twenty-six participants (63 females, ages 18-30 years; average: 20.87, STD: 2.56) with normal or corrected-to-normal vision and no reported neurological or psychiatric disease were recruited from the University of Maryland community. The project was approved by the University of Maryland College Park Institutional Review Board and all participants provided written informed consent before participation. Data from four participants were not used (one due to claustrophobia; one due to multiple episodes of falling asleep; and two given that the experiment was terminated before matching participants could be recruited (see below)).

### Anxiety questionnaires

Participants completed the trait portion of the Spielberger State-Trait Anxiety Inventory (STAI; Spielberger, Gorsuch et al. 1970) before scanning, and then completed the state portion immediately before the scanning session. The STAI scale is defined such that scores range from 20 to 80. In a recent analysis of over 3,000 individuals across behavioral and fMRI studies, the mode was a trait score of 36, and only 5% of the participants exhibited scores above 60 (Charpentier, Faulkner et al. 2020). Given our participant-matching procedure (see next), participants with trait anxiety scores above 60 were removed from the study.

### Participant matching

Participants completed the trait portion of the STAI during an initial screening interview. The earliest recruited participants were assigned to the controllable condition of the experiment. Each participant’s trait anxiety score and biological sex were used to find a match for the uncontrollable condition. In matching participants, we attempted to keep the difference in trait scores to within +/- 4 points. We prioritized matching new participants with already scanned participants. However, if we could not find a match with a +/- 4 point difference, we matched a new participant with participants who were already screened but not yet scanned. In such cases, we randomly assigned a participant to the controllable or uncontrollable condition. When necessary, matching of a new participant had to wait until a new one was screened who had an anxiety score within +/4 points. Overall, the time to recruit a match for controlled participants varied from one day up to one month.

### MRI data acquisition

Functional and structural MRI data were acquired using a 3T Siemens TRIO scanner with a 32-channel head coil. First, a high-resolution T2-weighted anatomical scan using Siemens’s SPACE sequence (0.8 mm isotropic) was collected. Subsequently, we collected functional EPI volumes using a multiband scanning sequence (Feinberg, Moeller et al. 2010) with TR = 1.0 sec, TE = 39 ms, FOV = 210 mm, and multiband factor = 6. Each volume contained 66 non-overlapping oblique slices oriented 30° clockwise relative to the AC-PC axis (2.2 mm isotropic). A high-resolution T1-weighted MPRAGE anatomical scan (0.8 mm isotropic) was collected. Additionally, in each session, double-echo field maps (TE1 = 4.92 ms, TE2 = 7.38 ms) were acquired with acquisition parameters matched to the functional data.

### Functional MRI preprocessing

We adopted the same procedures used in our previous study to minimize the impact of image distortion and improve spatial localization (see also Smith, Hur et al. 2018, Meyer, Padmala et al. 2019). To preprocess the functional and anatomical MRI data, we used a combination of packages and in-house scripts. The first three volumes of each functional run were discarded to account for equilibration effects. Slice-timing correction used the Analysis of Functional Neuroimages’ (AFNI; Cox 1996) 3dTshift with Fourier interpolation to align the onset times of every slice in a volume to the first acquisition slice. To reduce the contribution of head motion, we employed FSL’s Independent Component Analysis, Automatic Removal of Motion Artifacts (ICA-AROMA) (Pruim, Mennes et al. 2015). Components classified as head motion are regressed out of the functional MRI data with FSL’s fsl_regfilt.

In this study, we strived to improve functional-to-anatomical co-registration given the small size of some of the structures of interest. Skull stripping determines which voxels are considered part of the brain and plays an important role in successful subsequent co-registration and normalization steps. Currently, available packages perform sub-optimally in specific cases, and mistakes in the brain-to-skull segmentation can be easily identified. Accordingly, to skull strip the T1 high-resolution anatomical image (which was rotated to match the oblique plane of the functional data with AFNI’s 3dWarp), we employed six different packages [ANTs (Avants, Tustison et al. 2009; http://stnava.github.io/ANTs/), AFNI (Cox 1996; http://afni.nimh.nih.gov/), ROBEX (Iglesias, Liu et al. 2011; https://www.nitrc.org/projects/robex), FSL (Smith, Jenkinson et al. 2004; http://fsl.fmrib.ox.ac.uk/fsl/fslwiki/), SPM (http://www.fil.ion.ucl.ac.uk/spm/), and BrainSuite (Shattuck and Leahy 2002; http://brainsuite.org/)] and employed a “voting scheme” as follows (Smith, Hur et al. 2018, Meyer, Padmala et al. 2019): based on T1 data, a voxel was considered to be part of the brain if 4/6 packages estimated it to be a brain voxel; otherwise the voxel was not considered to be brain tissue.

Subsequently, FSL was used to process field map images and create a phase-distortion map for each participant (by using bet and fsl_prepare_fieldmap). FSL’s epi_reg was then used to apply boundary-based co-registration to align the unwarped mean volume registered EPI image with the skull-stripped anatomical image (T1 or T2), along with simultaneous EPI distortion-correction (Greve and Fischl 2009).

Next, ANTS was used to estimate a nonlinear transformation that mapped the skull-stripped anatomical image (T1 or T2) to the skull-stripped MNI152 template (interpolated to 1-mm isotropic voxels). Finally, ANTS combined the nonlinear transformations from co-registration/unwarping (from mapping mean functional EPI image to the anatomical T1 or T2) and normalization (from mapping T1 or T2 to the MNI template) into a single transformation that was applied to map volume-registered functional volumes to standard space (interpolated to 2-mm isotropic voxels). In this process, ANTS also utilized the field maps to simultaneously minimize EPI distortion.

The resulting spatially normalized functional data were blurred using a 4mm full-width half-maximum (FWHM) Gaussian filter. Spatial smoothing was restricted to gray-matter mask voxels. Finally, the intensity of each voxel was normalized to a mean of 100 (separately for each run). Intensity normalization was performed in the same manner on the non-smoothed version of the spatially normalized functional data.

### Voxelwise analysis

The voxelwise analysis followed the same model described in equation (1). Thus, the analyses were the same, with the exception of the multilevel component (ROIs). The cluster extent for statistical thresholding was determined by simulations using the 3dClustSim program and other AFNI tools. For these simulations, the smoothness of the data was estimated using 3dFWHMx (restricted to gray matter voxels) based on the residual time series from the participant-level voxelwise analysis. Taking into account the recent report of increased false-positive rates linked to the assumption of Gaussian spatial autocorrelation in fMRI data (Eklund, Nichols et al. 2016), we used the –acf (i.e., autocorrelation function) option added to the 3dFWHMx and 3dClustSim tools, which models spatial noise as a mixture of Gaussian plus monoexponential distributions. This improvement was shown to control false-positive rates around the desired alpha level, especially with relatively stringent voxel-level uncorrected p values such as 0.001 (Cox, Chen et al. 2017). Based on a voxel-level uncorrected p value of 0.001, simulations indicated a minimum cluster extent of 13 voxels for a cluster-level corrected alpha of 0.05.

### Visualization of response shape

Estimation of stressor responses relied on simultaneously accounting for all signal contributions, including fMRI responses during approach and retreat periods surrounding the stressor. To do so, response magnitude estimation of stressor events relied on convolving regressors with a canonical hemodynamic response. In doing so, regression coefficients are efficiently estimated for all conditions, allowing us to estimate responses to stressors. But to aid visualization of the stressor response, we performed an additional unassumed-shape analysis (also called deconvolution analysis) that estimated signal intensities at every time point following stressor delivery for a window of 13.75 seconds. We stress that the objective of this analysis was to help visualize the responses, and not to draw statistical inferences.

Responses were recovered using the 3dDeconvolve AFNI program using cubic spline basis functions, which provide a slightly smoother approximation of the underlying signal than using delta (“stick”) functions (also called finite impulse responses or FIR). The results (Figures 4 and 7) confirm that the canonical hemodynamic response provided an adequate response model for most brain regions, including the anticipated peak around 5 seconds post onset. However, deconvolution indicated that the canonical model was probably not ideal for some regions, such as the anterior hippocampus, PCC, and PCC/precuneus, which appeared to follow a different time evolution (note that the mismatch is not because the response was negative, but related to the shape/timing of the response itself).

Finally, we note that we chose to perform the main inferential analyses (described in the preceding sections) by using regressors convolved with the hemodynamic response because we did not have a priori information concerning the timing/shape of stressor-related responses. Importantly, stressor duration varied to some extent for each event, which poses a problem for deconvolution (essentially, each trial’s response varied because of timing differences and potentially due to “noise” fluctuations).

### Skin conductance response (SCR) analysis

For the analysis of SCR, we employed methods previously adopted in our past work (e.g., Meyer, Padmala et al. 2019). Each participant’s SCR data were initially smoothed with a median-filter over 50 samples (200 ms) to reduce scanner-induced noise. Previous work has capitalized on the slow evolution of SCR signals to propose an analysis approach paralleling the one used with fMRI (Bach et al. (2009) and Engelmann et al. (2015)). Accordingly, the pre-processed SCR data were analyzed using multiple linear regression using the 3dDeconvolve program in AFNI. We employed the same regression model as the one used for fMRI data. All regressors were convolved with a canonical skin conductance response model based on the sigmoid-exponential function (Lim, Rennie et al. 1997).

### Relationship between SCR and brain activity: Trial-by-trial analysis

To probe the relationship between brain activity and physiological arousal, we focused on the BST and the left dorsal anterior insula (see Results). Our goal was to evaluate the relationship between trial-by-trial responses in the brain and SCR. To estimate single-trial responses of fMRI data, the model was the same one employed for the participant-level analysis, except for the stressor regressor, which was defined separately for each stressor event (“trial”). Because the simultaneous presence of all trial-level stressor regressors in the model would unduly increase collinearity, estimation was performed via the iterative procedure developed by Mumford et al. (2012) as implemented by the 3dLSS program of the AFNI suite.

The same approach was employed to estimate trial-by-trial SCRs. General methods for SCR processing followed our past work (e.g., Meyer, Padmala et al. 2019). For computational tractability, SCR data were resampled by decimating the original 250 Hz sample rate to 10 Hz. Note that given the slow evolution of this type of signal, the downsampled time series preserved the information needed for the analysis. As with fMRI data, 3dLSS was employed to estimate single-trial responses.

### Bayesian multilevel model

The basic linear model can be written as

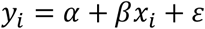

where *x* is a predictor variable. A multilevel model with a grouping/clustering variable (for example, ROIs) indexed by *j*, can be expresses as

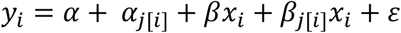

where *α* is the overall intercept, *β* is the overall slope, *α*_*j*[*i*]_ are *j* intercepts, and *β*_*j*[*i*]_ are *j* slopes (say, one per ROI). With two regressors, we can write

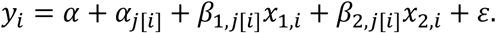

In the present study, the difference of stressor response, Δ_*p,r*_, for a yoked pair of participants *p* and ROI *r* can be expressed in terms of the linear mixed effects model notation (`lme4`-like) as

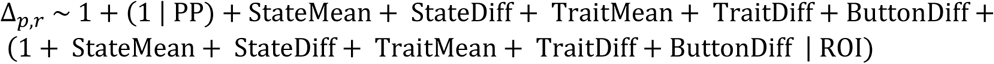

where PP (participant pair) and ROI are grouping variables, and the other terms indicate the covariates in the model. The first line specifies the intercepts, with a general one, as well as one per participant pair; it also specifies the 5 covariates (slopes). The second line specifies slopes for the covariates nested within ROI. The notation “1 + covariate” indicates that they are allowed to have a nonzero correlation with the intercepts (technically specified via a variance/covariance matrix).

The full model is therefore (in the notation of McElreath 2020):

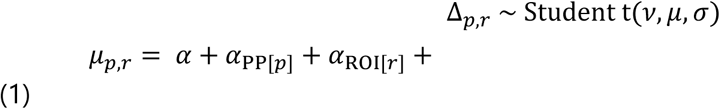

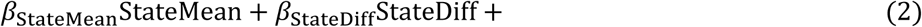

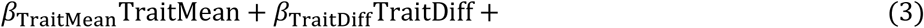

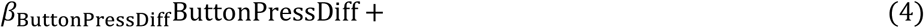

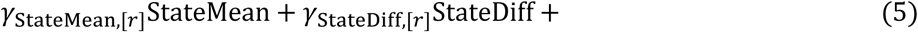

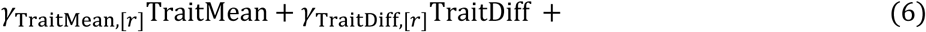

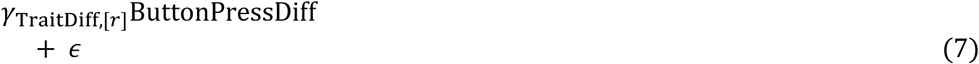

where the *α* terms are “intercepts”, with the “varying intercepts” *α*_*p*_ and *α*_*r*_ capturing the contribution of each participant pair *p* and each ROI *r*. The slope parameters *β* are parameters that model the contributions of the covariates, and the “varying slopes”, *γ*_*r*_, model the covariate contribution at each specific ROI.

In addition, the following weakly-informative priors were employed. All intercept and slopes were defined as

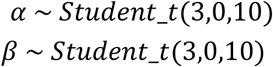

for all five covariates. The notation for the Student t distribution, *Student_t*(*v, μ, σ*), is such that *v* is the degrees of freedom, *μ* is the mean, and *σ* is the scale parameter. This prior is a very noninformative heavy-tail distribution suggested by Stan.

The intercepts clustered by participant were defined as

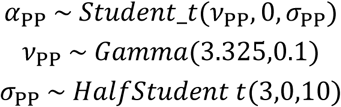

where the Gamma distribution, *Gamma*(*k, β*), is such that *k* is the shape parameter and *β* is the rate parameter. Note that *α*_PP_ expresses a vector of intercept parameters, one for each participant pair. The default suggested by Stan is *Gamma*(2,0.1). However, we chose our parameters because they provide 50% probability mass that a value can be above/below 30. As the Gamma prior informs the estimate of the degrees of freedom, a value of *v* > 30 essentially entails a normal distribution. Values less than 30 will amount to heavier tails and more skew/outlier tolerance. In any case, the actual estimates of *v* are estimated from the data, and the Gamma prior is rather weakly informative. The *HalfStudent_t*(*v, μ, σ*) is simply a positively truncated Student t.

In a similar fashion, the slopes clustered by ROI were defined as

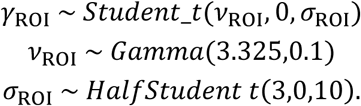

Finally, for the variance-covariance structure, the LKJ correlation prior (Lewandowski et al., 2009) was used with the shape parameter with the value 1.

## Supplementary tables

**Table 1.** Uncontrolled stressor vs. Controlled stressor

| Table 1 |  | Uncontrolled stressor vs. Controlled stressor |  |  |
| --- | --- | --- | --- | --- |
| Hemisphere | Area | MNI coordinates (X Y Z) | Cluster extent (voxels) | t-value at peak coordinate |
| *Uncontrolled > Controlled* |  |  |  |  |
| L | Middle/ inferior occipital gyrus | -46 -84 -2 | 17 | 4.100 |
| R | Middle occipital gyrus | 30 -78 16 | 24 | 4.523 |
| R | Fusiform gyrus (1) | 28 -74 -12 | 14 | 4.141 |
| R | Fusiform gyrus (2) | 36 -68 -12 | 16 | 4.385 |
| L | Inferior temporal/ occipital gyrus | -48 -68 -10 | 182 | 5.700 |
| L | Posterior intraparietal sulcus | -24 -68 34 | 60 | 4.550 |
| L | Superior parietal lobule | -24 -66 56 | 29 | 4.022 |
| R | Superior parietal lobule | 24 -64 48 | 46 | 4.750 |
| L | Anterior intraparietal sulcus | -46 -38 40 | 129 | 5.066 |
| L | Middle frontal gyrus/ precentral gyrus ( | -28 -4 58 | 177 | 4.886 |
| L | Inferior frontal gyrus/ precentral gyrus | -40 2 34 | 215 | 6.187 |
| L | Dorsomedial prefrontal cortex | -8 14 50 | 212 | 6.177 |
| L | Anterior insula | -38 20 -4 | 81 | 5.038 |
| R | Middle cingulate cortex | 8 20 46 | 14 | 4.609 |
| L | Anterior cingulate cortex | -6 36 30 | 14 | 4.647 |
| L | Inferior frontal gyrus | -48 46 0 | 17 | 3.740 |
| R | Cerebellum (lobule VIIb) | 30 -70 -50 | 25 | 4.193 |
| R | Cerebellum (lobule VI) | 34 -62 -28 | 43 | 4.744 |
| L | Putamen | -18 10 2 | 39 | 5.170 |
| L | Caudate | -14 14 8 | 45 | 4.781 |
| *Controlled > Uncontrolled* |  |  |  |  |
| R | Posterior cingulate cortex | 12 -56 18 | 49 | -4.802 |

Activation clusters defined based on voxel-level 0.001 threshold, minimum 13 voxel extent: Cluster corrected 0.05.
Abbreviations: L, left; R, right

**Table 2.** Exploratory Analysis

| Table 2 |  | Exploratory Analysis |  |  |
| --- | --- | --- | --- | --- |
| Hemisphere | Area | MNI coordinates (X Y Z) | Cluster extent (voxels) | t-value at peak coordinate |
| *Controlled > Uncontrolled* |  |  |  |  |
| R | Angular gyrus | 44 -74 34 | 25 | -3.721 |
| L | Supramarginal gyrus | -44 -36 26 | 17 | -4.304 |
| R | Transverse temporal gyrus | 44 -24 16 | 16 | -3.637 |
| L | Parietal operculum | -40 -18 20 | 12 | -3.396 |
| R | Superior temporal gyrus | 64 -20 -6 | 24 | -4.707 |
| R | Medial frontal gyrus | 18 30 44 | 12 | -3.943 |
| L | Medial frontal gyrus | -6 56 26 | 30 | -4.330 |
| L | Putamen | -28 -4 -10 | 10 | -3.942 |

Activation clusters defined based on voxel-level 0.005 threshold, minimum 10 voxel extent. Note that for a voxelwise threshold of 0.005, a cluster extent of 27 or more voxels is corrected at the cluster level at 0.05. Only Controlled > Uncontrolled reported.
A cluster in the posterior cingulate cortex is not reported here as it was included in Table 1.

## Supplementary figures

**Figure 1**: Supplementary Movie 1.

**Supplementary Figure 1.**
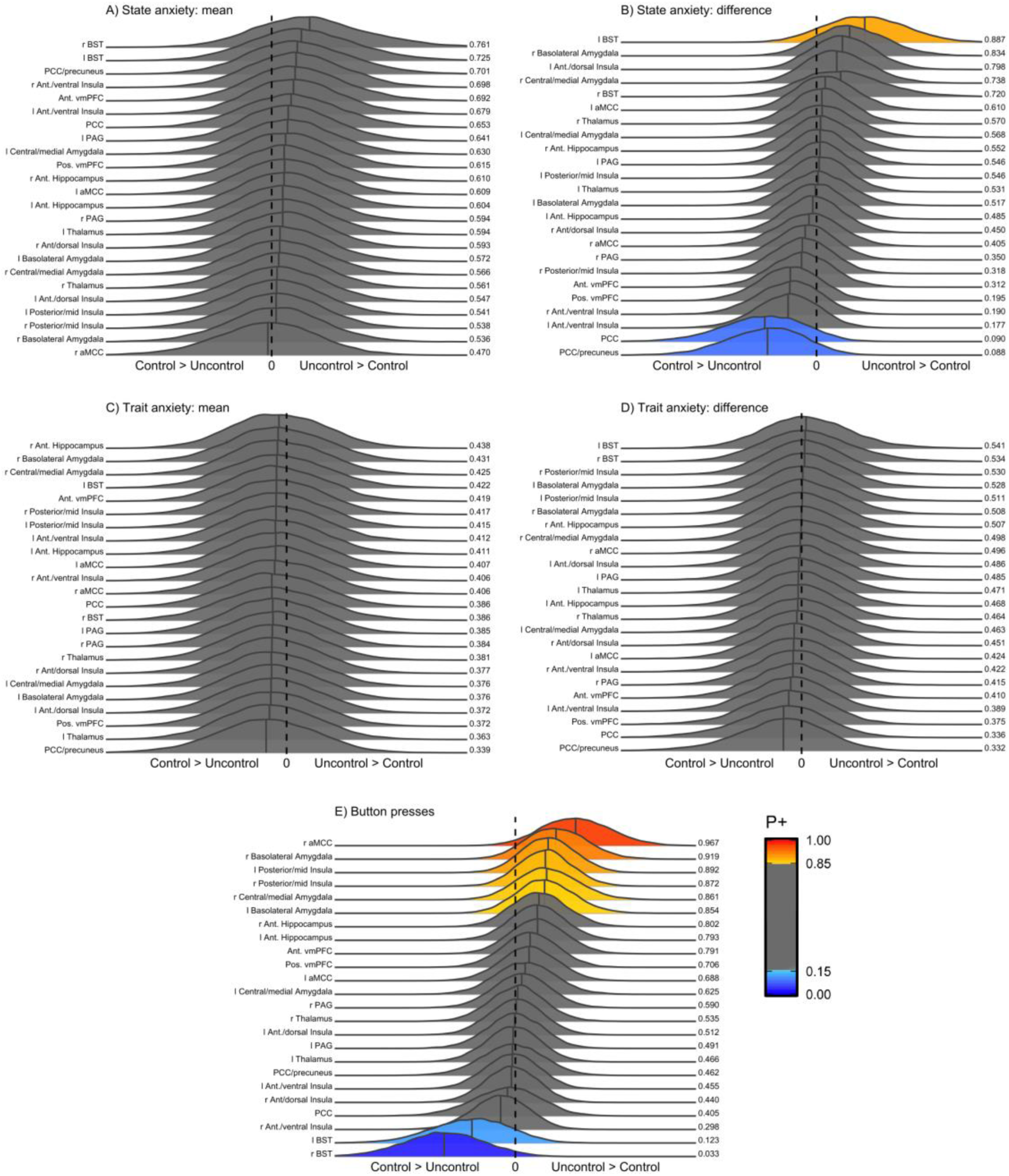
Posterior distributions for the five covariates. P+ indicates the probability that the effect is > 0.

**Supplementary Figure 2.**
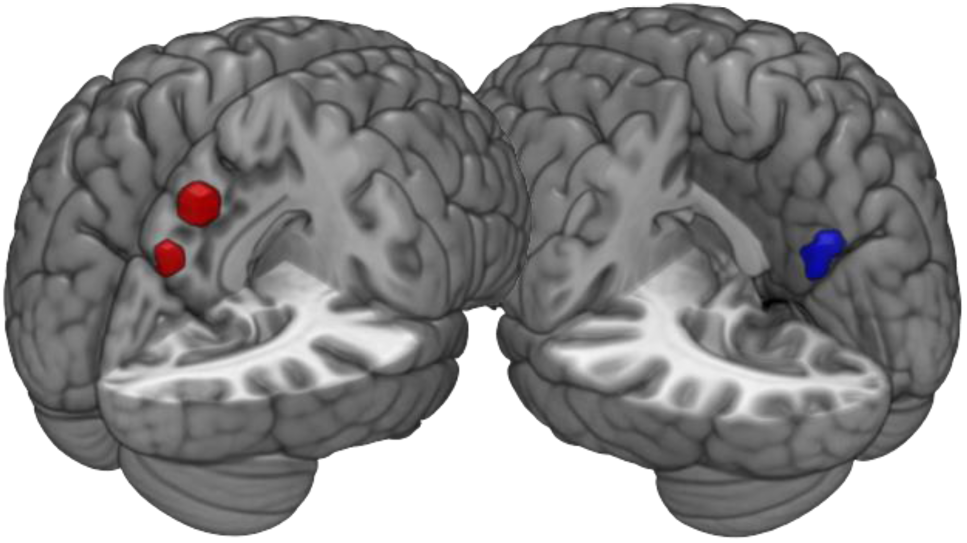
Left: PCC/precuneus and PCC ROIs. Right: activation cluster with a controllability effects (voxelwise analysis).

**Supplementary Figure 3.**
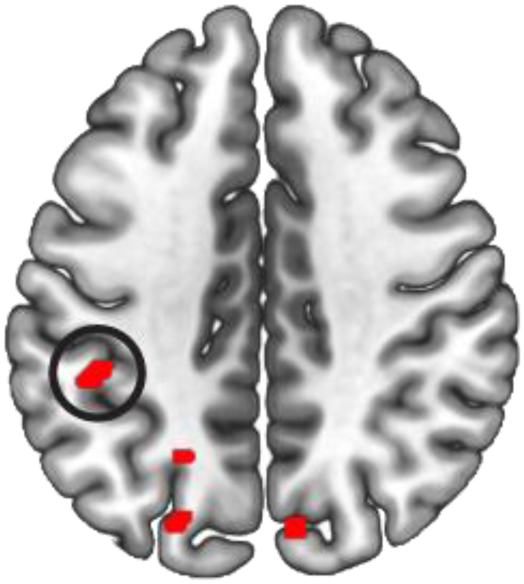
Activation map for the effect of button-press difference. The only cluster that overlapped the controllability effect is encircled. Thresholded at 0.01 voxelwise and 5 voxel cluster extent.

